# The non-specific Lipid Transfer Protein (nsLTP) is involved at early and late stages of symbiosis between *Alnus glutinosa* and *Frankia alni*

**DOI:** 10.1101/2021.10.29.465983

**Authors:** Mélanie Gasser, Nicole Alloisio, Pascale Fournier, Severine Balmand, Ons Kharrat, Joris Tulumello, Abdelaziz Heddi, Pedro Da Silva, Philippe Normand, Hasna Boubakri, Petar Pujic

## Abstract

*Alnus glutinosa* response to *Frankia alni* is driven by several sequential physiological modifications that include calcium spiking, root hair deformation, penetration, induction of primordium, formation and growth of nodule. Here, we have conducted a transcriptomic study to analyse plant responses to *Frankia alni* at early stages of symbiosis establishment.

Forty-two genes were significantly activated by either with a *Frankia* culture supernatant or with living cells separated from the roots by a dialysis membrane permitted to identify plant genes which expression changes upon early contact with *Frankia*. Most of these genes encode biological processes, including oxidative stress and response to stimuli. The most upregulated gene is the non-specific lipid transfer protein (nsLTP) encoding gene with a fold change of 141. Physiological experiments showed that nsLTP increases *Frankia* nitrogen fixation at sub-lethal concentration. Immunohistochemistry experiments conducted at an early infection stage indicated that nsLTP protein is localized at the deformed root hair region after *Frankia* inoculation and later in nodules, precisely around bacterial vesicles. Taken together, these results suggest that nsLTP acts at early and late stages of symbiosis, probably by increasing nitrogen uptake by *Frankia*.

## Introduction

Symbiosis between *Alnus glutinosa* and the actinobacterium *Frankia alni* permits the establishment of root nodules in which nitrogen fixation takes place. This helps alder and other plants collectively called “actinorhizal” to grow on nitrogen-poor soils and initiate ecological successions. The determinants of this interaction are poorly known, due among other reasons to the lack of a transformation system in alder. *Alnus glutinosa* is the type species of the genus (Navarro *et al*., 2003), it grows in pioneer biotopes such as glacial moraines, volcanic ashes and mine spoils (Normand & Fernandez, 2019) and its genome has been deciphered recently (Griesmann *et al*., 2018).

The presence of a symbiotic bacterium like *Frankia* or simply that of its exudates, triggers in the host plant tissues a series a events leading to root hair deformation and calcium spiking (Granqvist *et al*., 2015). Later, penetration, primordium formation and growth of nodules follow with subsequent exchange of nutrients. Omic analyses have permitted to gain a global view of the physiological changes taking place upon specific challenges. In particular, transcriptomics of 21 day post-inoculation (dpi) nodules permitted to see on the plant side the presence of homologs of the whole common symbiotic signalling cascade or CSSP (Hocher, V. *et al*., 2011; Hocher, Valérie *et al*., 2011), some of which have been lost in non-symbiotic neighbours (Griesmann *et al*., 2018).

This important conservation on the plant side of the common symbiotic pathway (Hocher, V. *et al*., 2011) contrasts with the absence in most *Frankia* genomes (Normand *et al*., 2007) of the canonical *nod* genes which are involved in the biosynthesis of Nod Factor (NF) in majority of Rhizobia (D’Haeze & Holsters, 2002). In mature nodules, the bacterial genes *nif, hup, shc* and *suf* are upregulated but no trace of a symbiotic island was found (Alloisio *et al*., 2010). A study of the *Frankia alni* early response (2.5 dpi) showed that several determinants were upregulated among which a K-transporter, lipid modifying enzymes and a conserved cellulose synthase cluster (Pujic *et al*., 2019). Proteomics was performed on a related species, *Frankia coriariae* that unravelled an upregulation of cell wall remodelling enzymes, signal transduction and host signal processing proteins (Ktari *et al*., 2017). Metabolomics on an *Alnus*-infective strain showed the presence of various compounds absent from roots (Hay *et al*., 2017), high levels of TCA intermediates citrate, fumarate and malate (Carro *et al*., 2015a) and high levels of citrulline, glutamate and pyruvate (Brooks & Benson, 2016; Hay *et al*., 2020). Signalling between alder and *Frankia alni* involves on the part of the bacterium the synthesis of an uncharacterized root hair deforming factor (Ghelue *et al*., 1997; Cérémonie *et al*., 1999) and that of the auxin PAA (Hammad *et al*., 2003). On the plant side, less is known besides the upregulation of defensins that modify the porosity of *Frankia* membranes (Carro *et al*., 2015b; Carro *et al*., 2016).

We undertook the present study to better understand the initial steps of the actinorhizal symbiosis on the plant side where the symbiont must be recognized to pave the way for its internalization. To analyse this early plant responses to *Frankia* symbiont we have conducted a cell-free contact through two conditions: indirect contact with *Frankia* trapped in dialysis tube or its supernatant and by targeting 2.5 dpi response, by which time extensive root hair deformation has occurred (Berry & Torrey, 1983) and before primordium formation at 7dpi (Lalonde, 1979). We further focussed on the most upregulated gene coding a non-specific Lipid Transfer Protein (nsLTP), which is also one of the most overexpressed genes in the nodule. We showed that nsLTP increases *Frankia* nitrogen fixation at sub-lethal concentration. Taken into account that this protein shown to localize at the deformed root hairs and in *Frankia’s* vesicles inside nodules, we hypothesized that nsLTP acts at early and late stages of symbiosis.

## Materials and methods

### Strains and growth condition before total RNA extraction

Plant samples at early stages of infection were obtained as described previously(Pujic *et al*., 2019). Briefly, *Frankia alni* ACN14a (Normand & Lalonde, 1982) was grown in BAP-PCM media until log-phase, collected by centrifugation, washed twice with sterile ultra-pure water and suspended in Farhaeus medium without KN03. *Frankia* cells were homogenised by forced passage through a series of needles (21G, 23G, 25G, 27G) before inoculation.

*Alnus glutinosa* seeds obtained from a single individual growing on the banks of the river Rhone in Lyon were surface sterilised and grown as described earlier (Pujic *et al*., 2019). Seedlings were transferred to Fåhraeus’ solution (Fahraeus, 1957) in opaque plastic pots (8 seedlings/ pot) and grown for four weeks with 0.5 g.L^−1^ KNO_3_, followed by one week without KNO_3_ before inoculation.

Two independent experiments were performed. In experiment #1 (Exp 1-*Frankia* indirect contact (*F*IC)), 8ml of *Frankia* cells were transferred to dialysis tubing (MWCO=100kDa) and arranged into a plastic pot containing 8 seedlings and filled with Fåhraeus’ solution without KNO_3_. Five biological replicates were performed with 4 plastic pots per replicate. In experiment #2 (Exp2-*Frankia* supernatant direct contact (SupC)), *Frankia* supernatants were applied directly on *Alnus* roots. Supernatant extract was prepared as follow: 250 ml log-phase culture cells were removed by centrifugation and the supernatant filtered through 0.22 µm membranes (Millipore, Billerica, MA). Solid phase extractions of the supernatants were done using benzene sulphonic acid cation exchanger on silica (Macherey Nagel, Hoerdt, France). Elution was done using 50:50 v/v methanol-water followed by 100 % methanol. Two fractions were lyophilised under vacuum, dissolved in water and tested for root hair deformation activity on independent plant roots. In parallel, as a control condition without supernatant, an equivalent volume of culture medium was treated with the same extraction protocol. Three biological replicates were performed with 6 seedlings per replicate.

The root hair deformation process took place similarly in both direct and indirect contact conditions, confirmed with stereomicroscope observations (Leica MZ8, Wetzlar, Germany). After 2.5 days (64 hours) with root hairs highly deformed, a 2 cm long segment representing the central part of the root was cut at 2 cm from the distal end, washed with sterile water before freezing in liquid nitrogen and storing at -80° C. In addition, a T0 control condition without *Frankia* was performed and plant roots were washed and frozen as described above.

### Transcriptomics of alder at 2.5dpi

Transcriptomics was done as described earlier (Hocher, V. *et al*., 2011). Total RNA was purified from roots using RNeasy plant mini kits (Qiagen, Courtaboeuf, France) and treated with DNases as before. Residual DNA was removed using the Turbo DNA free kit (Ambion, Thermo Fisher Scientific, Wilmington, DE), quantified using a NanoDrop spectrophotometer (Thermo Fischer Scientific) and qualitatively assessed using a Bioanalyzer 2100 (Agilent, Waldbronn, Germany).

Microarrays were designed (13909 probes; 1 probe/*A. glutinosa* unigene), manufactured and hybridised by Imaxio (http://www.imaxio.com/index.php), using Agilent Technologies (http://www.home.agilent.com/agilent/home.jspx) as previously described (Hocher, V. *et al*., 2011). The microarrays were scanned with an Agilent G2505C Scanner. The Feature Extraction software (Agilent, version 11.5.1.1) was used to quantify the intensity of fluorescence signals and microarray raw data were analysed using GeneSpring GX 12.0 software (Agilent technologies). Normalisation per chip (to the 75th percentile) and per probe (to the median) were performed to allow comparison of samples. The two experiments Exp 1 and Exp 2 were analysed separately. In order to limit false positive results, microarray data were filtered according to the flag parameter “detected”. Thus, probes taken into account are uniform, non-outlying, non-saturated and displayed an intensity level above the background in at least one of the two biological conditions.

In Exp1, 1590 probes yielding an intensity level significantly below the background in the 10 samples were discarded. In addition, 1146 probes did not yield the flag “detected” in at least one of the two biological conditions and were also discarded. Thus, 11 173 probes were taken into account for subsequent analyses of Exp 1.

In Exp 2, 1765 probes yielded an intensity level significantly below the background in the 6 samples and were discarded. In addition, 1113 probes did not yield the flag “detected” in at least one of the two biological conditions and were also discarded. Thus, 11031 probes were taken into account for subsequent analyses of Exp 2. T-tests comparing *F*IC roots *vs*. control roots in Exp 1 and SupC roots *vs*. control roots in Exp 2 were applied and only those genes with an average fold change (FC) above 2 (up-regulated) or below 0.5 (down-regulated) with a *p*-value<0.05 were considered significant. The normalized and raw microarray data values have been deposited in the Gene Expression Omnibus database (www.ncbi.nlm.nih.gov/geo; accession nos. E-MTAB-8936 and E-MTAB-8937).

### Quantitative real-time RT-PCR

Reverse transcription (RT) and real time quantitative PCR (qRT-PCR) were performed with the same biological replicates used for microarray experiments. For *A. glutinosa* analyses, RT was performed with 5 µg of total mRNA using Transcriptor Reverse Transcriptase and oligo (dT)_15_ primer (Roche, Mannheim, Germany). QRT-PCR was run on a LightCycler 480 (Roche) using LightCycler 480 SYBR Green I Master (Roche) under the following conditions: 95 °C for 5 min; 45 cycles of 95 °C for 20 s, 60 °C for 20 s and 72 °C for 15 s. Primer sets were designed using ProbeFinder (Roche) and Primer 3 softwares and can be seen as Supporting Information (Table S1). Two qRT-PCR reactions were run for each biological replicate and each primer set. Expression values were normalised using the expression level of the *Ag-ubi* gene that encodes ubiquitin (Hocher, V. *et al*., 2011).

### Cloning of *agLTP24* gene

The *agltp24* gene was PCR amplified from cDNA prepared above using designed primers (Table S1) under the following conditions: 98 °C for 30 s; 30 cycles of 98 °C for 7 s, 63 °C for 20 s and 72 °C for 8 s. The reaction mixture of 50 µl PCR contained 1X Phusion HF buffer, 200 µM dNTPs, 0.5 µM Forward primer, 0.5 µM reverse primer, 87 ng template DNA and 1-units Phusion DNA Polymerase (NEB, Evry, France). The PCR product was cloned using the hot fusion method (Fu *et al*., 2014) in pET30a+ vector (Merck, Molsheim France) pre-digested with *Eco*RI and *Bgl*II in order to fusion AgLTP24 with a N-terminal 6XHis flag. Ligation mixture was transformed in *Escherichia coli* DH5α chemocompetent cells (Chung *et al*., 1989) for plasmid DNA propagation and sequencing (pET30a-HIS-AgLTP24, supplemental Fig. S1). Plasmids DNA was prepared using (Nucleospin Plasmid DNA kit Macherey Nagel) from 4 independent clones, 2 ml of each *E. coli* culture grown overnight in 5ml LB medium at 37°C with shaking. Cloned insert DNA was checked by Sanger sequencing (Biofidal, Lyon, France) and used for electrotransformation into iSHuffle T7 *lysY E. coli* cells (NEB, Evry, France), a strain used to obtain proteins with disulphide bonds (Lobstein *et al*., 2012).

### AgLTP24 production and purification

Cultures were grown in LB medium supplemented with kanamycin (40µg.ml^−1^) at 30°C at 130 rpm until log phase was reached (OD_600_ ∼0,5) then the HIS-AgLTP24 expression was induced with 1 mM isopropyl β-D-1-thiogalactopyranoside (IPTG). Induced cultures were incubated at 30°C for 3h at 130 rpm and harvested by centrifugation (5100 g for 10 min at 4°C). Expression was analysed by 4-20% Tricine SDS-PAGE (Schagger, 2006). Pellets were lysed by freeze thaw and resuspended in a binding buffer (100 mM Tris, 500 mM NaCl, 5mM Imidazole, pH 7.4). A second lysis was performed using silica beads and Fastprep 24™ 5G (MP Biomedicals). The cell lysate was harvested by centrifugation at 5100 g, 4°C for 30min and used for the purification of HIS-AgLTP24 using His GravityTrap columns (GE Healthcare) with modified binding (100mM Tris, 500mM NaCl, 5mM Imidazole, pH 7.4) and elution buffers (100 mM Tris, 500 Nacl, 500 mM Imidazole, pH 7.4). Purification of HIS-AgLTP24 was verified by 4-20% Tricine SDS-PAGE. Elution fractions was applied to a 3K MWCO Amicon Ultra-15 centrifugal filter (Merck Millipore) to buffer exchange with the enterokinase buffer (20 mM Tris, 50 mM NaCl, 2 mM CaCl_2_, pH 6.8) and concentrate. The HIS tag was removed from the mature sequence of AgLTP24 with enterokinase (NEB, Evry, France) for 16 h at room temperature. Cleavage was confirmed by 4-20% Tricine SDS-PAGE. The enterokinase was removed from the mixture using the Enterokinase Removal Kit (Sigma). His tag removal was done with a His GravityTrap column (GE-Healthcare) using an imidazole free binding buffer (100 mM Tris, 500 mM NaCl, pH 7.4). Wash fractions containing AgLTP24 purified were applied to a 3K MWCO Amicon Ultra-15 centrifugal filter (Merck Millipore) for buffer exchange with NH_4_^+^-free FBM medium (FBM-). Final concentration was determined with Qubit Protein Assay Kit (ThermoFisher). The purified AgLTP24 protein were further analysed by UPLC-ESI-HRMS. The system used for UPLC-ESI-HRMS was an ultra-performance liquid chromatography Ultimate 3000 (Thermo Fisher Scientific, Villebon-sur-Yvette, France) coupled to a high resolution hybrid quadrupole-time of flight mass spectrometer (Impact II, Bruker, Brême, Germany) equipped with electrospray ionization source (ESI) (Bruker, Brême Germany). Instrument control and data collection were performed using Data Analysis 5.0 software.

### Immunolocalization of AgLTP24

Immunostaining and fluorescence microscopy was done as described earlier (Login *et al*., 2011) with the synthesised epitope (DKHSTADFEKLAPCGKAAQD) injected to a rabbit (Covalab). Plant immunolocalization was performed on *A. glutinosa* roots (2.5 dpi) as described above in Exp 1 (*F*indC). In addition, nodules (21dpi) obtained as a previous study (Carro *et al*., 2015b) were also used here to localise agLTP24 at the mature stage of symbiosis. T0 non-infected seedlings were used as control.

### Physiological assays with AgLTP24 on *Frankia alni*

The antimicrobial activity of AgLTP24 (using 1µM, 2µM, 5µM concentrations), was done using the resazurin assay in 24-well microplates (Greiner bio-one, Les Ulis, France). A 2-week-old *Frankia* ACN14a culture grown in FBM medium supplemented by 5mM of NH_4_Cl (FBM+) was centrifuged at 5100 g for 15 minutes, the supernatant was removed, and the pellet homogenized in FBM-by repeated passage through syringes. *Frankia* culture was done in 1ml of FBM-per well with a final optical density of 0.02 and with an appropriate concentration of AgLTP24 peptide. Four replicates were made for each condition. Negative (*Frankia* ACN14 in FBM-) and positive controls (*Frankia* ACN14a in FBM-supplemented by kanamycin at 40µg.ml^−1^ final concentration) were performed. Microplates were incubated at 28°C, 80 rpm in a humid incubator. After 6 days of incubation, resazurin (Fisher Scientific SAS, Illkirch, France) was added to a final concentration of 6.25µg.ml^−1^ in each well and the microplates were incubated at 28°C, 80 rpm for 16 hours in the dark. Fluorescence measurements were performed with a microplate reader (infinite M200PRO, TECAN) with an excitation wavelength of 530nm and an emission wavelength of 590nm. The results obtained were normalised to the mean fluorescence of the negative control. Since data were normally distributed, mean comparisons with the negative control were performed with a Student’s *t*-test.

A second bioassay was made by growing *F. alni* in FBM- and incubating from 1 to 14 days at 28°C (5 replicates by kinetic point and by condition) as described previously. At each kinetic point, the effects nitrogen fixation (or ARA), respiration (IRA) and on morphology were monitored (Prin *et al*., 1990; Carro *et al*., 2015b). Since data were normally distributed or not, different statistical tests were made using GraphPad Prism 9.2.0 (GraphPad software Inc; San Diego, CA, USA).

### Bioinformatics

Blast analyses and GO assignation were performed on Blast2GO v5.2 using BLASTx against NR, Gene Ontology (GOs) and Inter-ProScan (Conesa & Gotz, 2008). Fishers’ exact test implemented in Blast2GO was used to identify significantly enriched GO categories. A GO category was considered significantly enriched only when the *p*-value for that category was < 0.05 after applying FDR correction. Molecular mass of proteins and their isolectric point were calculated through the Expasy software (https://web.expasy.org/protparam/)

## Results

### Transcriptomics response of alder at 2dpi

In the first early response transcriptomic experiment (Exp 1), 300 and 225 genes were found to be significantly up- or down-regulated in *A. glutinosa* after 2.5 days post inoculation (dpi) of *Frankia* in a separated dialysis bag (cut off 100 KDa -*Frankia* indirect contact (*F*indC) roots/ control roots) by having a fold change (FC) ≥ 2 or ≤ 0.5 (p ≤ 0.05), respectively (Table S2). In Exp2, 248 *A. glutinosa* genes were found to be significantly regulated 2.5 dpi with *Frankia* supernatant (*Frankia* supernatant contact (*F*supC) roots/control roots). Of these, 176 and 76 genes were also up- and down-regulated in Exp2; respectively (Table S3). By comparing ESTs present in both experiments, 84 genes were found significantly up- or down-regulated (See SI Table S4).

### Biological processes implicated in early steps of symbiosis

A GO-based analysis of the biological processes enriched in up- and down-regulated genes within each experiment yielded 12 to 23 % of genes with no Blast hit (Fig. S2). Among the sequences with a Blast hit, 8 to 14% correspond to unknown proteins and could not be associated to any biological process. The annotated sequences retrieved were distributed in 21 different processes (Fig. **1a**). The presence of *Frankia* at early stages strongly activated genes involved in response to stimuli and catabolic processes among which oxidative stress (oxidase, peroxidase) and defence (chitinase, defensin, thaumatin…) were upregulated in both conditions, as well as a large portion of genes involved in nitrogen compound metabolic processes with kinases, transferases, lyases or glucosidases. Globally, the number of sequences retrieved was higher in Exp1 than Exp2 (Fig. **1a**), suggesting a stronger signal detected in indirect interaction through dialysis tubing.

**Fig. 1:**
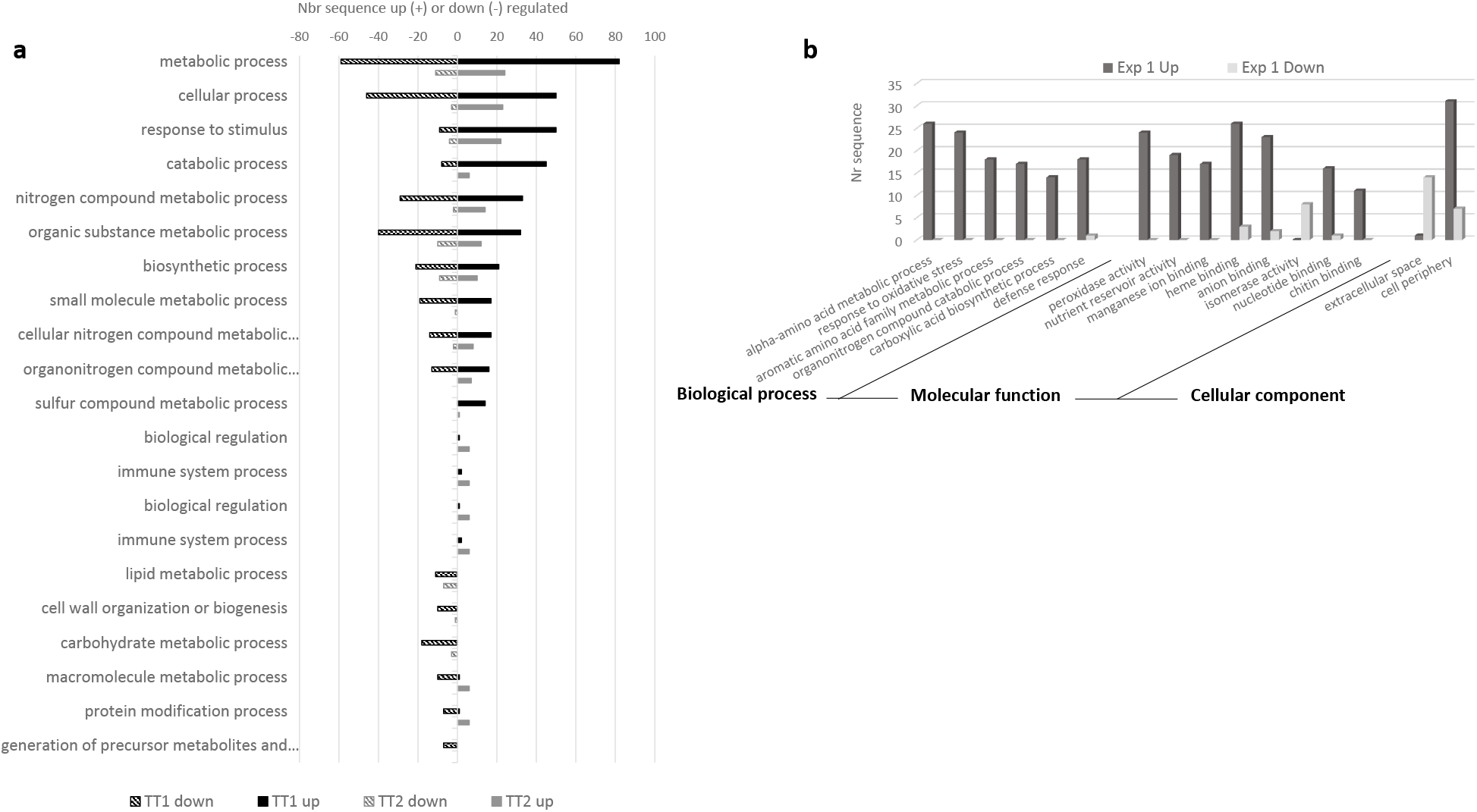
Gene Ontology (GO) functional classification analysis. **a**. Distribution of annotated genes into functional categories according to GO biological process. **b**. GO functional classification analysis was made by comparing up and down-regulated genes found in Exp1 (Table S2) according to biological process, molecular function and cellular component through Fisher test. Number of sequences (Nr) were indicated for each category. BLAST2GO pipeline was used to get this GO annotation.

A statistical analysis permitted to identify certain enzyme families by comparing each pool of genes. Even this Fisher test made in both experiment, only Exp1 gave differences in GO assignation between up and down-regulated genes (Fig. **1b**).

In Exp1, an overrepresentation of genes coding enzymes involved in oxidation-reduction biological processes and oxidative stress responses was seen with numerous peroxidases, catalases, oxidases or lipoxygenases overexpressed (Table S1). Also, chemical reactions involving alpha and aromatic amino acid metabolism and fatty acid beta-oxidation, selenocysteine methyltransferase, phenylalanine ammonia lyase or caffeic acid O-methyltransferase were seen following indirect contact with *Frankia* cells. Nucleotide binding proteins such as a receptor kinase (AG-N01f_037_C05), a uridine kinase (AG-N01f_030_B06) or a calcium transporting membrane (AGCL1701Contig1) were also more abundant in the set of up-regulated proteins of Exp1. Finally, several genes coding defence response were overrepresented such as endochitinase, disease resistance protein, defensin, thaumatin, allergen or nsLTP. Conversely, we observed an enrichment in down-regulated genes of one molecular function of GO assignation: isomerase such as inositol-3-phosphate synthase, ribose-5-phosphate isomerase or beta-amyrin synthase.

### The most upregulated genes in both experimentations

By compiling differentially regulated genes found in both experiments (Table 1 and Table S4), we compared them through GO-based analysis. An overrepresentation of only one biological process (lipid metabolism) in down-regulated genes was found with the same statistical restriction (FDR<0.05). Indeed, several genes, including synthase, reductase and oxidase and involved in triterpenoid (AG-N01f_010_H21) isopentenyl diphosphate (IPP) and dimethylallyl diphosphate (AG-N01f_014_J09); geranyl or geranylgeranyl diphosphate (GPP and GGPP) (AG-N01f_005_E08; AG-J07f_004_F15) or terpene (AG-J07f_004_M14; AGCL3086Contig1) biosynthesis were detected as more abundant in down-regulated genes and already observed by our first GO assignation (Fig. **1a**).

**Table 1:**
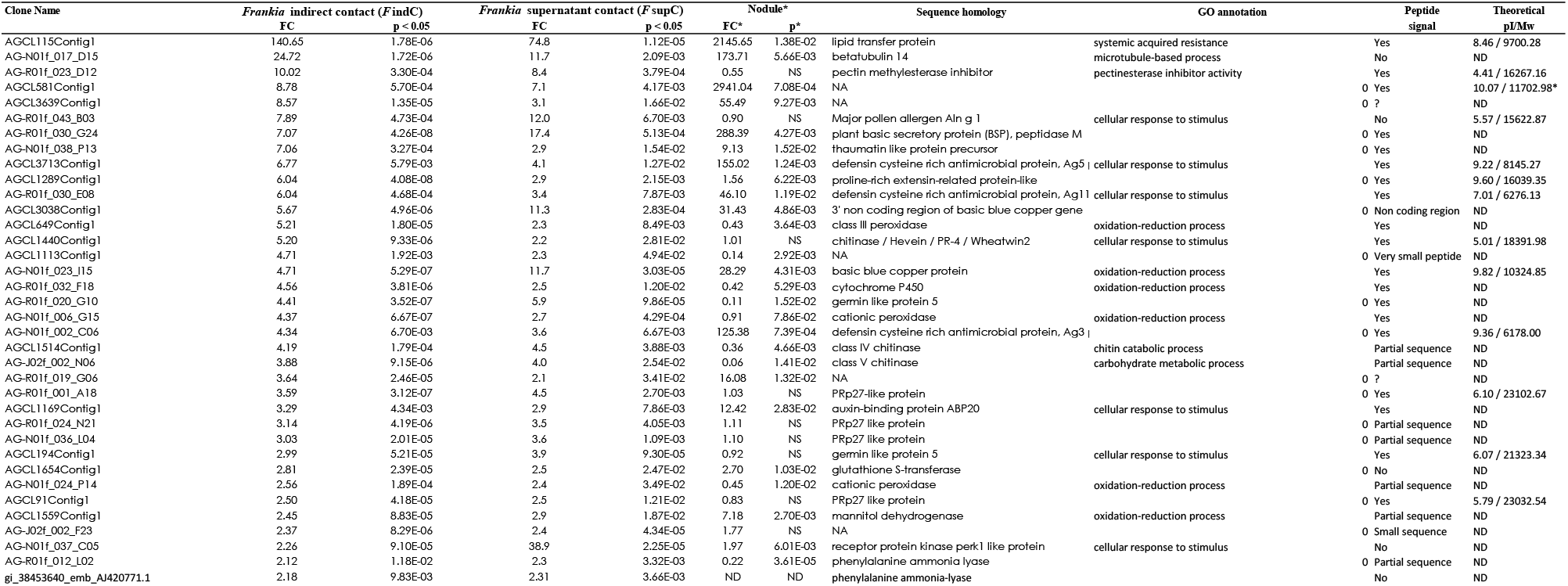
The up-regulated genes during early root hair deformation of *Alnus glutinosa* in both experiments, with *Frankia* indirect contact (*F* IndC) and with *Frankia* supernatant contact (*F* SupC) For contigs, **EMBL-Accession** numbers of all the contigated EST sequences are indicated. Singleton or contig **nucleotide sequences** are indicated when they were not of high-enough quality for EMBL submission acceptance. Fold Change (**FC**) is the ratio between *F* IndC roots and control roots and between *F* SupC roots and control roots, respectively. The significant difference of gene expression between *F* IndC roots and control roots and between *F* SupC roots and control roots (FC ≥ 2 or ≤ 0.05) was assessed by a T-test at p<0.05 **\***:Results of the previously reported microarray study with *Alnus glutinosa* nodules are indicated: **FC\*** is the ratio between nodules and non inoculated root and **p\*** is the p value of a T-student test comparing these two conditions (Hocher et al, 2011). ns indicated non significant difference between nodules and non inoculated root (p<0.05) For EMBL-accession number, see Suppl Table 3. ** AGCL3038Contig1 sequence is the 3’ non coding region of basic blue copper gene (AG-N01f_023_I15).

Amongst the 42 most upregulated genes found in both conditions (Table 1), the most upregulated gene encodes a non-specific lipid transfer protein or nsLTP (fold changes: 140.7 (*F*indC) and 74.8 (*F*supC)), followed by a betabutilin and a pectin methylesterase inhibitor. Remarkably, 20 out these 42 up-regulated genes encoded proteins with a peptide signal (Table 1). Among them, numerous upregulated genes encode secreted peptides (47%), some of them are classified as antimicrobial: three defensins (Ag5, Ag3 and Ag11), a nsLTP and a thaumatin like-protein.

To confirm the level of expression at this early step, we focussed on two genes, one coding the nsLTP and the second coding the basic blue copper protein (fold changes: (fold changes: 140.7 (*F*indC) and 74.8 (*F*supC), 4.7 (*F*indC) and 11.7 (*F*supC), respectively) as candidate genes for qRT-PCR with two primer pairs per gene (Table S1). Up-regulation was confirmed by qRT-PCR with the same biological replicates previously used (Table S5).

In order to examine the temporal distribution of expression, we extracted from our previous work (Hocher, V. *et al*., 2011) the fold change obtained with the same microarray but at 21 dpi when the nodule is formed and compared it to non-infected roots (Table 1, Fig. 2).

**Fig. 2:**
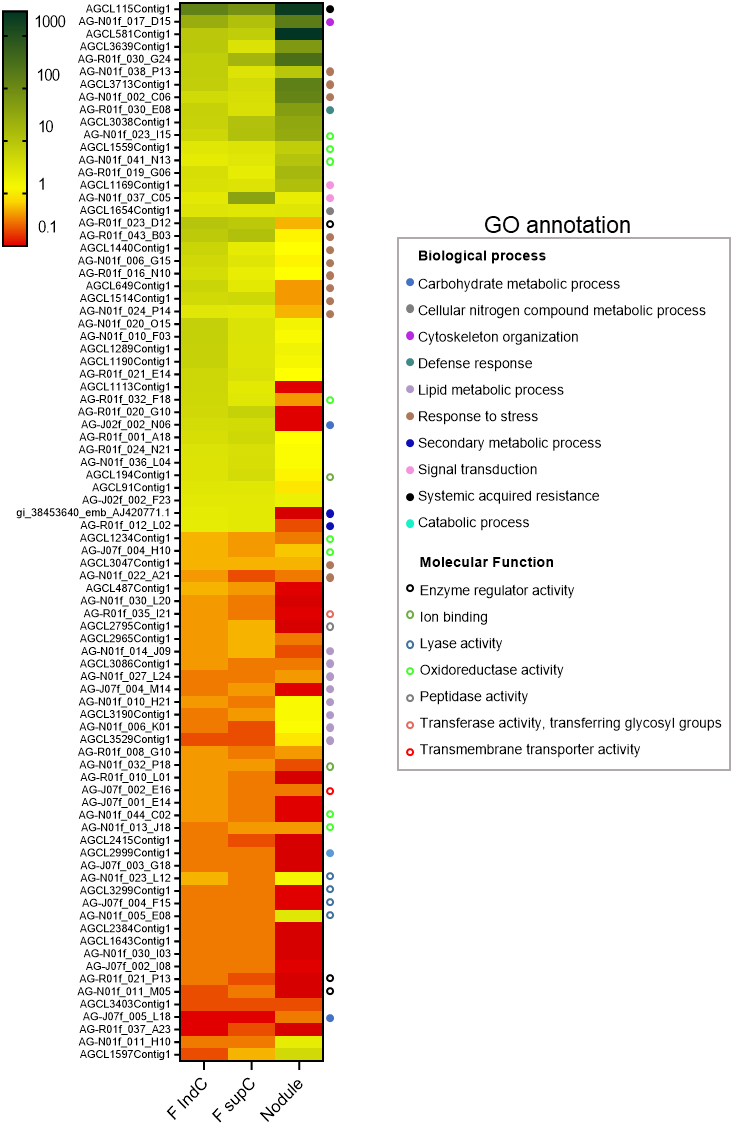
Expression levels of the genes which were commonly upregulated and downregulated in *F*indC and *F*supC conditions. The expression level values in nodule were extracted from our previous work ((5); Table S4). The expression values were expressed as Fold change (FC) with a color gradient from red (down-regulated; ≤0.5) and orange-yellow (no-regulated) to green (up-regulated; ≥2) in infected roots or nodules compared to control roots without *Frankia*. The GO annotations were assigned by noting preferentially biological process and if not present, the molecular function. Certain genes have no GO assignation after blast2Go analysis. The heatmap was generated by GraphPad Prism 9.2.0.

Almost down-regulated genes at early stage were either repressed or not modulated at nodule stage. Among them, we observed an enrichment of proteins associated to lipid metabolic process and lyase activity. Conversely, genes upregulated at 2.5 dpi are classified in three profiles. Sixteen are still induced when 17 are not modulated and 8 are repressed in nodule. For instance, genes encoding proteins involved in stress response are found in the three profiles (Fig. 2) but we found mainly AMPs (defensin, thaumatin) in the first group when chitinase and peroxidase enriched the two other groups (Table 1). The nsLTP found as the most upregulated gene at early stage presented also a very high expression at 21 dpi (FC=2145 in nodule, Table 1) suggesting a requirement all along the symbiosis process. NsLTPs are classified as AMPs and annotated as systematic acquired resistance according GO analysis suggest its potential role in defense response. For all these reasons, we decided to focus our functional analysis on this protein.

### Structure and classification of AgLTP24

The nsLTP from *A. glutinosa* overexpressed during its interaction with *Frankia* is named AgLTP24. The total nucleotide sequence is 552 base pairs and contains one exon. This sequence encodes a putative processed protein of 114 amino acids including a signal peptide with a sequence of 22 amino acids that allows its secretion [29]. The mature peptide has a molecular weight of 9700.28 Dalton and a cationic isoelectric point of 8.46, this nsLTPs is classified according to Edstam et al. (Edstam *et al*., 2011) as Type D. It contains the characteristic motif of nsLTPs composed of 8 cysteines: C-X13-C-X14CC-X9-C-X1-C-X24-C-X10-C which stabilises the 3D structure by folding the α-helices domains with disulphide bridges forming a hydrophobic cavity.

### Immunolocalization of AgLTP24 in planta

In order to determine where this nsLTP is secreted during symbiosis establishment, we performed an immunohistolocalization within plant tissues. Antibodies anti-AgLTP24 were applied to different alder tissues inoculated or not by *Frankia*. First, we produced deformed root hairs after an indirect contact of *Frankia* (F indC) and observed signals in extracellular nooks (Fig. **3b** and Fig. **3c**). These nooks were the specific site for *Frankia* binding. No signal was detected with control rabbit serum (Fig. **3a**) or with anti-AgLTP24 against non-infected roots (Fig. S3). Microarrays and qRT-PCR showed overexpression of this protein in mature 21 dpi nodules. We observed a specific immunolocalization in plant cells infected by *Frankia* situated in the fixation zone of the nodule (Fig. **3e**), specifically on *Frankia’*s vesicles (Fig. **3f**).

**Fig. 3:**
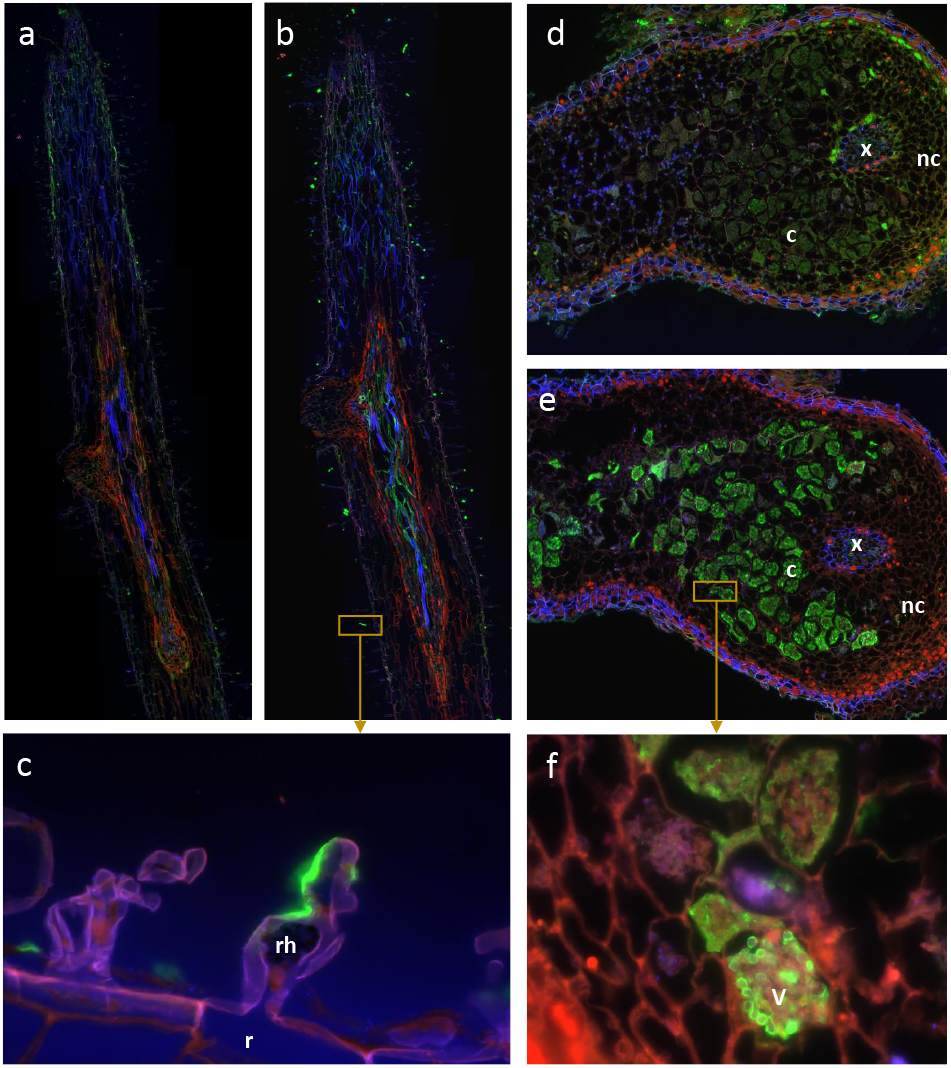
AgLTP24 localization in roots after 2.5 days (a, b, c) or nodule of 21 dpi (d, e, f). Hair magnification of a specific zone of the infected root **(c)** or nodule **(f)**. Immunofluorescence localization of AgLTP24 in hairy roots **(b, c)** and around *Frankia* vesicles in infected cells **(e, f)**. Negative control with rabbit serum before intramuscular injection of AgLTP24 epitope showed no green fluorescence in hairy roots **(a)** or infected cells **(d)**. Blue: DAPI; red: autofluorescence; green: Alexa Fluor 488 dye anti-rabbit antibody (Life Technologies, Saint Aubin, France). c, infected cells; nc, noninfected cells; v, *Frankia* vesicles; x, xylem; rh, root hairs.

### Biological production of AgLTP24

The nsLTPs are low molecular weight peptides rich in cysteines. It has a signal sequence that allows the plant to address it to a target compartment. Those nsLTPs are characterized by a conserved motif of 8 cysteines in their mature sequence forming 4 disulfide bonds important for the formation of the hydrophobic cavity, which makes their production difficult. Synthetic production is limited to small peptides with few disulfide bonds, so we developed a biological production of mature AgLTP24, i.e. without its signal peptide, in *E. coli* shuffle T7 *lys*Y (Fig. S1) which are capable of forming disulfide bonds (Lobstein *et al*., 2012). We obtained a yield of 0.1mg of pure protein per litre of culture. After peptide purification, UPLC-ESI-HRMS analysis of the purified AgLTP24 protein revealed a monoisotopic [M+H]^+^ m/z 9686.8008 (theoretical monoisotopic m/z calculated for C_415_H_676_N_123_O_126_S_9_: 9686.7760) consistent with the formation of four disulfide bridges (Fig. S4). This analysis was made after each run of purification and before bioassays to check the purity and the good conformation of the protein with disulphide bridges.

### Physiology of *Frankia alni* in contact with AgLTP24

A miniaturised, rapid and fluorescent bioassay using resazurin dye was successfully developed to determine whether AgLTP24 had antimicrobial activity against *Frankia alni* at 7 dpi. For this, we applied a range of concentrations of AgLTP24 from 1 to 5µM (Fig. 4) and used kanamycin as positive control of inhibition of *Frankia*. The cell viability of *Frankia* is reduced from a concentration of 5µM AgLTP24 (54 ± 16% of cell viability) while no effect was detected below this concentration. AgLTP24 can thus be considered an antibacterial peptide against the symbiotic partner at this concentration. In addition, microscopy observation showed a negative effect of AgLTP24 on vesicles’ production (Fig. **4b**). Indeed, at 5µM of AgLTP24, no vesicle was observed whereas they are present at sub-inhibitory concentrations. However, *Frankia* must be viable and efficient in the nodule to ensure nitrogen fixation for trophic exchange with its plant partner. This conducted us to perform a second physiological assay by using sub-inhibitory concentrations of AgLTP24. The second physiological assay was conducted with a range of AgLTP24 from 1nM to 1µM (Fig. 5). In this test, nitrogen fixation, respiratory activity (OD_490nm_) and growth effects (OD_600nm_) were monitored over a 10 days’ time course (Fig. S5).

**Fig. 4:**
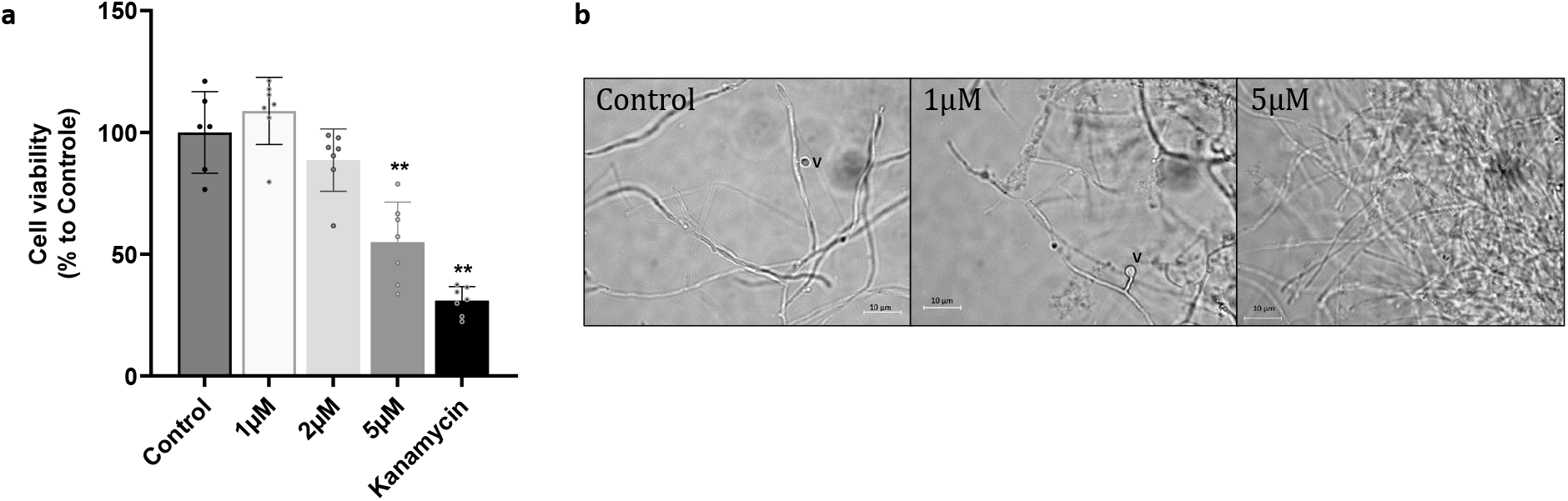
physiological bioassays on *Frankia* growth and cellular development. **a**. Resazurin bioassay on *Frankia* ACN14a growth after supplementation of 1 to 5µM of AgLTP24. Cell viability was calculated by normalizing the fluorescence of the assay with the mean of the negative control (*Frankia* without AgLTP24). Kanamycin (40µg.ml^−1^) was used as positive control. Data are expressed as mean values ± SD. Differences between normalized data were assessed by the Mann Whitney test (bilateral and unpaired) compared to control. Graphic representation and statistical analysis of results was conducted with GraphPad Prism version 9.2.0. *p value <0.05, **p value <0.01.**b**. *Frankia* cells observation without or with gradual concentration of AgLTP24 from 0 (Control) to 5µM and obtained during the different physiological bioassays. v, *Frankia* vesicles

**Fig. 5:**
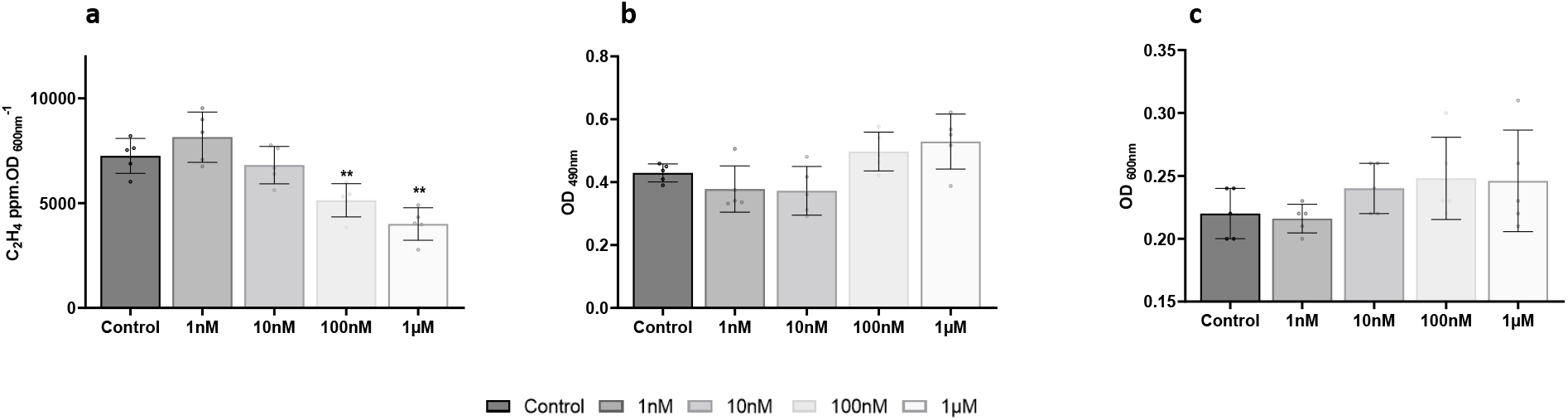
**effect of AgLTP24 in physiology of *Frankia*** after 7 days of growth on **a**. Nitrogen fixation; **b**. respiratory activity and **c**. growth by measuring OD_600nm_. Data were extracted from the kinetic assay (Fig. S5). Data are expressed as mean values ± SD. Differences between means were assessed by the Mann Whitney test (bilateral and unpaired) compared to control. Graphic representation and statistical analysis of results was conducted with GraphPad Prism 9.2.0. \**p* value <0.05, \*\**p* value < 0.01.

That range of AgLTP24 had little effect on *Frankia* growth. We noted a negative effect at 10nM of AgLTP24 on respiratory activity at early time of the time course (Fig. **S5b**) but it was not maintained over time. Regarding respiratory activity, we observed a positive but not significant effect with AgLTP24 above 100nM and 5 days of growth. The most striking effect was observed on nitrogen fixation after 7 days of culture (Fig. 5) with a concentration above 100nM leading to inhibitory effect whereas no effect was observed on growth or respiratory activity. Conversely, at 1nM of this peptide, the AgLTP24 improved this activity. Microscopy did show any effect on vesicle morphology (data not shown).

## Discussion

Early steps of the interaction between alder and *Frankia* start with the molecular dialogue through the secretion of flavonoids by the plant (Benoit & Berry, 1997; Hughes *et al*., 1999) and the bacterial “Nod-like” factors by *Frankia* (Cérémonie *et al*., 1999; Perrine-Walker *et al*., 2011). This factor also called root hair deforming factor (RHDF) because it triggers plant response by reorientating root-hair tip growth followed by the formation of an infection thread (IT) and cell division induction in inner cortical cells. This process leads to a nodule where *Frankia* will be housed and exchange with its plant partner. Even though RHDF structure remains unknown, previous studies have shown that this factor of around 3KDa is present in *Frankia* cell free supernatant and acts at nanomolar dose (Cissoko *et al*., 2018). In addition, this factor is able to induce a high frequency nuclear Ca^2+^ spiking (Granqvist *et al*., 2015). Due to the lack of a genetic transformation system for alder, other approaches besides genetic inactivation have been used to identify the host symbiotic determinants.

Focusing at nodule step (Hocher, V. *et al*., 2011), transcriptomic analysis showed that the common symbiotic cascade known in Legumes (interacting with rhizobia) and in most land plants (interacting with VAM fungi) (Genre & Russo, 2016) was also present in actinorhizal plants (Gherbi *et al*., 2008; Hocher, V. *et al*., 2011; Hocher, Valérie *et al*., 2011). Previously, differential screening of cDNA libraries from root and nodules of *Alnus glutinosa* with nodule and root cDNAs permitted to identify several symbiotic genes among which a subtilisin-like protease (Ribeiro *et al*., 1995), a sucrose synthase, an enolase (van Ghelue *et al*., 1996), and a dicarboxylate transporter (Jeong *et al*., 2004). These genes are evocative of tissue reorganization and trophic exchanges but shed no light on the triggering of organogenesis. The present study aimed at investigating the plant molecular response at early symbiotic stages by using same transcriptomic microarray as in our previous work at nodule step (Hocher, V. *et al*., 2011).

### Global plant response at early step of symbiosis

Firstly, the two experiments induced different level of response in the plant partners with stronger signal in the Exp1, suggesting that even though the supernatant is sufficient to trigger root-hair deformation, a sustained dynamic of interaction with *Frankia* is more efficient. These results suggest that *Frankia* secretome produced during Exp1 could be more diverse triggering a strong perception of its presence by plant. Indeed, *Frankia* secretes a variety of proteolytic enzymes such as glycosidases, esterases, or proteases presumably involved in root infection (Mastronunzio *et al*., 2008).

Plant global response reveals a common pattern with a strong modulation of genes involved in stress response. This biological process, specific to early response (Hocher, V. *et al*., 2011) is constituted by genes encoding putative proteins involved in oxidative stress (oxidase, peroxidase); to defense against pathogens such as pathogenesis related (PR) proteins (chitinase, Defensin, thaumatin or nsLTP). Beside these biological processes, we can propose different steps in the plant response after *Frankia* contact.

#### Oxidative stress in hairy roots repressed in nodule

Focusing on oxidative stress, both the overall GO analysis showed a strong induction of genes involved in oxido-reduction metabolism (oxidase, peroxidase). These enzymes involved in oxidative stress to reduce reactive oxygen species (ROS) suggesting that *Frankia* supernatant induces a ROS stress such as previously observed for rhizobia at early steps of interaction (Peleg-Grossman *et al*., 2009; Peleg-Grossman *et al*., 2012). In legume roots, the enzymatic activities of catalase, ascorbate peroxidase, glutathione reductase or NADPH oxidase significantly increase upon inoculation with bacterial symbiont (Bueno *et al*., 2001; Den Herder *et al*., 2007; Peleg-Grossman *et al*., 2009; Peleg-Grossman *et al*., 2012). After this important release, ROS production reduced drastically to prevent root hair curling and IT formation (Shaw & Long, 2003; Lohar *et al*., 2007).

To follow this dynamics of expression over the course of symbiosis establishment, we extracted transcriptional data from our previous work made at nodule step (Hocher, V. *et al*., 2011). Few genes are not modulated or repressed in the nodule steps (Table S4, Fig. **2**), suggesting a similar profile. Consequently, plant after *Frankia* internalization would repress this stress response but the measure of ROS abundance over the time course of the symbiosis is necessary to support this tendency.

#### Perception of Frankia factor, signaling and hormonal function

Little is known about the symbiotic signals produced by *Frankia* but their perception requires plant receptor. Here, a statistical analysis revealed a moderate overrepresentation of gene involved in signal transduction (Table 1; AG-N01f_037_C05) encoding a putative receptor like kinase (RLK) somewhat repressed in the nodule suggesting its potential implication in *Frankia* recognition. However, other RLK similar to Lys6/Lys7/Nfr1 of legume NF receptors (AG-R01f_025_F02) already found in *A. glutinosa* (Hocher, V. *et al*., 2011) are not modulated after 2.5 days, which raises the question of *Frankia* recognition. Plant RLKs have been shown to control the initiation, development, and maintenance of symbioses with beneficial mycorrhizal fungi and rhizobia (Buendia *et al*., 2018; Chiu & Paszkowski, 2020) but the mechanism in actinorhizal symbiosis remain unknown (Svistoonoff *et al*., 2014).

The CSSP transduction will trigger primordium organogenesis where hormonal balance plays a crucial function and particularly auxin. Indeed, exogenous auxins treatments lead to the formation of thick lateral roots resembling nodules in actinorhizal Fagales (Hammad *et al*., 2003; Svistoonoff *et al*., 2003) and an auxin influx inhibitor perturbs the formation of nodules in another actinorhizal model, *Casuarina glauca* (Peret *et al*., 2008). Transcriptional data at early steps sustain the pivotal function of auxin by demonstrating the overexpression of several genes encoding auxin binding proteins (Table 1: AGCL1169Contig1) or auxin inducing proteins (Table S2: AGCL1376Contig1; AG-N01f_043_C20). We did not detected genes involved in salicylic acid (SA) or jasmonic acid (JA) biosynthesis in early steps of symbiosis however a fine bioinformatics study of metabolic routes using the genome alder is required to facilitate putative assignation and to validate or not their absence in those omic data.

In any case, perception of pathogen by plant triggers SA and JA accumulation and leads to the accumulation of PR proteins to minimize pathogen load or disease onset. In legume symbiosis, those hormones inhibit bacterial infection and nodule development (Liu *et al*., 2018) suggesting that NF recognition represses this accumulation.

#### Defense mechanisms activation

After oxidative stress, following *Frankia* contact, alder activates numerous genes encoding PR proteins which are key components of plant innate immune system against both biotic and abiotic stresses (Ali *et al*., 2018). More precisely, PR proteins have diverse functions such as glucanase, chitinases, thaumatin like, peroxidases, defensins, nsLTPs or thionins.

We found numerous upregulated chitinases as observed in legumes (Staehelin *et al*., 1994; Goormachtig *et al*., 1998; Xie *et al*., 1999) and described as important for rhizobia infection and IT formation (Malolepszy *et al*., 2018). Also, genes for oxidative stress and chitinases are not modulated or repressed in late step of actinorhizal symbiosis (Table S4, Fig. **2**) suggesting they are important in the early steps of *Frankia* infection. Furthermore, genes encoding one uncharacterized PR proteins is upregulated after *Frankia* contact (AG-R01f_016_N10) and rapidly repressed at nodule step (Table 1). A similar profile was observed for PR10 in *Medicago trunculata* when associated to symbiont or pathogen infection, it is activated but in symbiotic pathway, its expression is transitory suggesting that defense signaling pathways are suppressed during the establishment of symbiosis (Chen *et al*., 2017). Here, the transitory activation of the Alder PR protein could be also induced by *Frankia* factors even though a specific response must be demonstrated by comparing to other biotic and abiotic stresses.

Some of the PR proteins are classified as AMPs based on their small size, the conservation of cysteine rich motif and their potential action as antimicrobial compound (Tam *et al*., 2015). In alder transcriptome, we distinguished the overexpression of gene encoding plant defensins (Ag3, Ag5 and Ag11), thaumatins (AG-N01f_038_P13) and nsLTPs (AGCL115Contig1). Here, all AMPs induced at early steps are still overexpressed in nodule (Table S4, Fig. **2**).

Like defensin, thaumatin like proteins considered as a member of defense protein but also in plant development (de Jesús-Pires *et al*., 2020) is overinduced in alder during actinorhizal symbiosis. This expression profile is quite different to the well-documented thaumatin like gene in soybean. Indeed, the *rj4* thaumatin like gene was described as constitutively and similarly transcribed in roots and nodules and intervene at a very early stage of IT formation to inhibit nodule formation with an incompatible strain (Hayashi *et al*., 2014). This suggests that thaumatin-like proteins in alder could intervene in incompatibility function but its overexpression in nodule step also raise question about another function in nodule organogenesis.

### AgLTP24: from *Frankia* infection to nodule functioning

#### Function at early step

Global omic approach made at early steps of alder symbiosis pointed to a strong signal from one protein belonging to nsLTPs family named here AgLTP24. The immunolocalization on curled hairy roots induced after *Frankia* contact showed the specific binding of AgLTP24 in nooks (Fig. **3**) where *Frankia* embedding before internalization through IT formation. This suggests that AgLTP24 acts at the early phase of infection. This is supported the induction of an nsLTP (MtN5) in *M. trunculata* at early stage of symbiosis and in the nodule (Pii *et al*., 2009; Pii *et al*., 2012). Because its silencing resulted in an increased number of curling events with a reduced number of invading primordia, whereas its overexpression resulted in an increased number of nodules, authors concluded that MtN5 is important at very early step of infection for the successful establishment of the symbiotic interaction but not in nodule formation (Pii *et al*., 2013). Similar to MtN5 (Pii *et al*., 2009), AgLTP24 possesses a slight antimicrobial activity *in vitro* against its symbiont at 5µM. As *Frankia* cell integrity must be preserved for infection, we hypothesize that AgLTP24 concentration in contact with *Frankia* must be below to 5µM. Further investigation to assess protein concentration in plant tissues is a great challenge but requires the development of high-performance mass spectrometry technology coupled to imagery (Gemperline *et al*., 2015; Gemperline *et al*., 2016). Finally, AgLTP24 could be a gene player at early step of symbiosis to transduce signal within plant roots to permit IT formation as well as induce stress response in *Frankia* cells.

#### Function at late step

*AgLTP24* is also present in nodule (Table 1) and targets specifically *Frankia’s* vesicle in nodule fixation zone (Fig. **3f**) but this localization is different from MtN5 binding in the distal zone of the nodule (Pii *et al*., 2012). However, a nsLTP (AsE246) found in *Astragalus sinicus* specifically symbiosome membranes through binding a lipid component: digalactosyldiacylglycerol (DGG) (Lei *et al*., 2014). In this model, the *asE246* silencing impacts nodule development with fewer matured infected plant cells. Thus, AgLTP24 could fix vesicle wall composed by a multilamellate hopanoid lipid envelopes (Berry *et al*., 1993; Nalin *et al*., 2000) or plant perisymbiontic membrane coating *Frankia* cells in nodule. The perisymbiosome membrane composition is less documented but non-inoculated *Alnus rubra* roots are mainly composed of glycerol, phosphatidylethanolamine, phosphatidylserine and phosphatidylcholine (Berry *et al*., 1991). A lipidomic study in *A. glutinosa* roots using recent technology followed by a bioassay binding is a great perspective to decipher AgLTP24 mechanisms involved. As observed *in vitro*, AgLTP24 could act as antimicrobial compound by reducing cell viability and vesicles formation of *Frankia* (Fig. 4) suggesting a potential role by controlling *Frankia* proliferation in nodule. However, because expression of nitrogen fixation gene cluster is more active in symbiotic conditions compared to N-free culture condition (Alloisio *et al*., 2010; Lurthy *et al*., 2018), the concentration probably present in the nodule fixation zone would be lower to this lethal concentration. It is worth to note that at 1nM concentration of AgLTP24, the nitrogen fixation activity is induced a higher but not statistically so significant without perturbing *Frankia* growth (Fig. 5). The improvement of nitrogen fixation was also observed for the AgDef5 defensin translocated by *A. glutinosa* to *Frankia*’s vesicle in nodule (Carro *et al*., 2015b).

In addition, this defensin is also upregulated at 2.5 dpi (Table 1) suggesting the plant could deliver a cocktail of molecules including AgLTP24 to improve bacterial infection and nitrogen fixation in nodule. A depth investigation of this synergic effect *in vitro* opens a new perspective in addition to their genetic silencing in plant as an exciting challenge to complete the gap of knowledge regarding their role in actinorhizal symbiosis.

## Acknowledgements

Thanks are expressed to Elise Lacroix and the Greenhouse facility (FR 41), to Danis Abrouk in the ibio platform for help in submission of omic data to international database, Jonathan Gervaix in the AME platform to access ARA measurement and Philippe Bulet for his technical advice on peptide purification. We thank the PGE platform for other measurements. This project has been funded by the FR BioEnviS (Biodiversité, Environnement et Santé, Université Lyon1, France) and the EC2CO (Ecosphère Continentale et Côtière) grant (reference: 10459).

## Author contribution

M.G., N.A., A.H., P.N., P.P. and H.B. conceived and designed the study. M.G., N.A., P.F., S.B., O.K, J.T., P.D.S., P.P and H.B carried out the experiments. M.G., P.N., P.P. and H.B. performed the data analysis. M.G., P.D.S., and H.B. performed the figure drawing. M.G., P.N., P.P., A.H. and H.B. provided critical biological interpretations of the data. P.N. and H.B. wrote the manuscript.

